# Myeloid Cell Derived IL1β Contributes to Pulmonary Vascular Remodeling in Heart Failure with Preserved Ejection Fraction

**DOI:** 10.1101/2023.05.18.541302

**Authors:** Vineet Agrawal, Jonathan A. Kropski, Jason J. Gokey, Elizabeth Kobeck, Matthew Murphy, Katherine T. Murray, Niki L. Fortune, Christy S. Moore, David F. Meoli, Ken Monahan, Yan Ru Su, Thomas Blackwell, Deepak K. Gupta, Megha H. Talati, Santhi Gladson, Erica J. Carrier, James D. West, Anna R. Hemnes

## Abstract

**Background:** Pulmonary hypertension (PH) in heart failure with preserved ejection fraction (HFpEF) is a common and highly morbid syndrome, but mechanisms driving PH-HFpEF are not well understood. We sought to determine whether a well-accepted murine model of HFpEF also displays features of PH in HFpEF, and we sought to identify pathways that might drive early remodeling of the pulmonary vasculature in HFpEF.

**Methods:** Eight week old male and female C57/BL6J mice were given either L-NAME and high fat diet (HFD) or control water/diet for 2,5, and 12 weeks. Bulk RNA sequencing and single cell RNA sequencing was performed to identify early and cell-specific pathways that might regulate pulmonary vascular remodeling in PH-HFpEF. Finally, clodronate liposome and IL1β antibody treatments were utilized to deplete macrophages or IL1β, respectively, to assess their impact on pulmonary vascular remodeling in HFpEF.

**Results:** Mice given L-NAME/HFD developed PH, small vessel muscularization, and right heart dysfunction after 2 weeks of treatment. Inflammation-related gene ontologies were over-represented in bulk RNA sequencing analysis of whole lungs, with an increase in CD68+ cells in both murine and human PH-HFpEF lungs. Cytokine profiling of mouse lung and plasma showed an increase in IL1β, which was confirmed in plasma from patients with HFpEF. Single cell sequencing of mouse lungs also showed an increase in M1-like, pro-inflammatory populations of Ccr2+ monocytes and macrophages, and transcript expression of IL1β was primarily restricted to myeloid-type cells. Finally, clodronate liposome treatment prevented the development of PH in L-NAME/HFD treated mice, and IL1β antibody treatment also attenuated PH in L-NAME/HFD treated mice.

**Conclusions:** Our study demonstrated that a well-accepted model of HFpEF recapitulates features of pulmonary vascular remodeling commonly seen in patients with HFpEF, and we identified myeloid cell derived IL1β as an important contributor to PH in HFpEF.

## INTRODUCTION

Pulmonary hypertension is a heterogenous syndrome defined by invasively measured mean pulmonary arterial pressure > 20 mmHg. The most common worldwide cause of PH is due to left heart failure, and the subset of patients with heart failure with preserved ejection fraction and pulmonary hypertension (PH-HFpEF) carry the worst prognosis.^1^ Despite studies demonstrating similar degrees of vascular remodeling between patients with PH-HFpEF and other conditions such as pulmonary arterial hypertension (PAH),^2^ a limited understanding of the pathogenesis of PH-HFpEF has precluded the development of therapies that improve morbidity and mortality in PH-HFpEF. A key limiting factor in advancing our understanding of the pathogenesis underlying PH-HFpEF has been the relative paucity of animal models that faithfully recapitulate human features of PH-HFpEF.^3^ Models of HFpEF with concomitant PH have been reported in the literature in larger animals or rodents, but relatively fewer murine models of HFpEF have been reported in the literature that mimic features of pulmonary vascular remodeling seen in patients.^3, 4^

One widely accepted multi-hit murine model that recapitulates many of the cardiac and extra-cardiac features of the cardiometabolic subtype of HFpEF uses a combination of metabolic and hemodynamic stress induced by high fat diet and N(gamma)-nitro-L-arginine methyl ester (L-NAME), however the pulmonary vascular changes in this model are thus far little studied.^5^ In this study, we sought to determine the extent to which this model could recapitulate the pulmonary vascular remodeling phenotype commonly associated with HFpEF in humans,^4, 6^ and we further sought to determine pathways that might drive pulmonary vascular remodeling. Our study demonstrates that the combination of L-NAME and high fat diet also recapitulates pulmonary vascular remodeling features of HFpEF. Our findings, corroborated through human samples as well as a second animal model of HFpEF, suggest that myeloid cell derived IL1β is an important contributor to adverse pulmonary vascular remodeling in HFpEF.

## Methods

All animal studies were approved by the Vanderbilt University Institutional Animal Care and Use Committee (IACUC). Collection of human plasma samples and tissue sections from human autopsies was approved by the Institutional Review Board at Vanderbilt University Medical Center.

### Animal Studies

For all studies, male and female C57/BL6J mice were purchased at 8 weeks of age from Jackson Laboratories (Bar Harbor, ME, USA). Male and female db/db mice were purchased from Jackson Laboratories at the age of 16 weeks of age (Bar Harbor, ME, USA). To induce the HFpEF phenotype, C57/BL6J mice at 8 weeks of age (Jackson Laboratories, Bar Harbor, ME) were administrated L-NAME in sterile water (0.5 g/L) (pH 7.0) and given a 60% lipid content high fat diet ad libitum for 2, 5, or 12 weeks (BioServ, Flemington, NJ, USA). Mice in the control group received regular water with a calorie neutral 6% lipid content low fat diet (BioServ, Flemington, NJ). At each time point, mice underwent anesthetized echocardiography, open chest catheterization, and tissue harvest as previously described and in Supplementary Methods.^6^ Structural analysis of lung tissue included hematoxylin and eosin stain as well as immunostain for muscularization of vessels as previously described.^7^ Cytokine levels were measured by pooling plasma and lung lysate from 5 mice treated with either L-NAME/HFD or control water/diet for 5 weeks and was assayed using an antibody-based cytokine array (R&D Systems, ARY006) and was completed according to manufacturer’s instructions. Mouse IL1β in plasma was measured by ELISA (Thermo, BMS6002). Details of human studies are outlined in Supplemental Methods.

### Monocyte/Macrophage and IL1β Depletion Studies

To deplete circulating monocytes and phagocytic cells, C57/BL6 were given twice weekly intraperitoneal injections of 100 μl of 5 mg/ml of either clodronate liposomes or phosphate buffered containing liposomes (Liposoma, Amsterdam, Netherlands; Batch Nos: C29E0622, P20E0522). Mice were concurrently treated with either L-NAME/HFD or control water/diet, as outlined above, for 5 weeks. To deplete IL1β, C57/BL5 mice were either first treated with L-NAME/HFD or control for 3 weeks. During weeks 4 and 5, they were given twice weekly intraperitoneal injections of 100 μg of either IL1β (BE0246, BioXCell, Lebanon, NH) or isotype control antibody (BE0091, BioXCell, Lebanon, NH). Treatment in the final two weeks was chosen as a “reversal” treatment strategy as well as to prevent the confounding factor of development of autoantibodies to the antibody treatments.

### Bulk RNA Sequencing

Whole lung was flash frozen after treating C57/BL6 mice with either 5 weeks of either L-NAME/HFD or control water/diet. RNA was isolated using the RNEasy kit (Qiagen) and submitted to Novogene (Sacramento, CA) for paired end 150 sequencing on an Illumina platform. A nominal read depth of 30 million RNA (60 million ends) per animal was used.

These were uploaded to and analyzed on the Partek platform, using the STAR aligner to align to the mm39 reference mouse genome. An average of 96% of reads aligned to genome (94%-97%). Counts were normalized to Counts Per Million.

### Single cell RNA-sequencing

Mouse lungs were cleared of red-blood cells by cardiac perfusion with 3mL of sterile saline, then were excised and placed in a C-Tube (Miltenyi) (one C-tube/mouse) containing 5mL lung digest solution (5mL phenol free DMEM (Gibco), Dispase (Roche 04942078001), collagenase (Sigma-Aldrich C0130-1G), and 5uL DNAse (10,000iU). Lungs were dissociated using a gentleMacs dissociator for 17 minutes at 37 degrees Celsius. Cell suspensions were passed through 100 μm then 70 μm cell strainers and washed 3-times in 5mL cell buffer (phenol free DMEM, 0.2mM EDTA, and 0.5% FBS). Cells were centrifuged at 500g for 10 minutes. Red blood cells were removed by lysing with 2mL ACK buffer (KD Medical RGC-3015) for 5 minutes. Cells were then washed, centrifuged, passed through a 40 μm cell strainer and resuspended for counting. Cell-hashing was performed by incubating 1X10^6^ cells with Total-SeqB (B0301-B0304) cell-hashing antibodies (Biolegend) for 30 minutes. Cells were then washed 3 times in cell buffer, and equal numbers of cells from each of the 4 mice per condition were pooled into a single reaction for scRNA-seq library generation using the 10X Genomics Chromium 3’v3 kit. Library sequencing was performed on an Illumina Novaseq6000 targeting 50,000 reads per cell.

### scRNA-seq analysis

Alignment and demultiplexing were performed using CellRanger v7.0.1. Ambient RNA filtering was performed using CellBender v0.2.2.^7^ ScRNA-seq analysis was performed using Seurat^8^ v5 adapted from our previous work^9–11^ with additional data visualization using scCustomize (https://doi.org/10.5281/zenodo.7534950). Demultiplexing was performed using the HTOdemux function, and singlet cells were carried forward for downstream analysis. Following quality-control filtering (excluding cells with <500 or >5000 genes, and cells with >10% mitochondrial reads), data were normalized and scaled using the SCTransform function, including mitochondrial and ribosomal percentages as regression variables, followed by principal components analysis, neighborhood graph calculation and UMAP embedding using 45 dimensions, and recursive louvain-clustering/subclustering. Heterotypic doublet clusters were identified by co-expression of lineage-specific marker genes, and cell annotation was performed manually, informed by published reference datasets.^12, 13^ Cell-type specific differential expression was performed on SCT-transformed counts using the Wilcoxon test. Gene ontology enrichment analysis was performed using PANTHER.

### Statistical Methods

Physiologic findings from animal studies are presented as mean ± standard error (SEM). Data were analyzed in GraphPad Prism using one-way analysis of variance (ANOVA) with Tukey’s multiple comparisons. Group differences in bulk RNA sequencing data were assessed using Kruskal-Wallis test, and gene ontology determined using Webgestalt (webgestalt.org). In single cell sequencing analysis, cell-type specific differential expression was performed on SCT-transformed counts using the Wilcoxon test. Gene ontology enrichment analysis was performed using PANTHER.

### Data Availability

All animal data are available upon request to the corresponding author. Data sets will be deposited in the Gene Expression Omnibus (GEO) database at NCBI (accession numbers pending). Single cell sequencing raw genomic data will be available through GEO (accession numbers pending). Code used for scRNA-seq analysis and figure generation will be available at https://github.com/kropskilab/myeloid_il1b.

## RESULTS

### L-NAME and High Fat Diet Causes Pulmonary Vascular Remodeling

We first confirmed that treatment with L-NAME and high fat resulted in a phenotype consistent with HFpEF. Mice treated with L-NAME/HFD demonstrated preserved cardiac output and left ventricular ejection fraction, increased systemic blood pressure, increased left atrial diameter, and increased weight gain (**Suppl Figs 1,2**), all consistent with features of HFpEF. In mice treated with L-NAME/HFD, catheterization of the right ventricle also demonstrated an increase in right ventricular systolic pressure as early as 2 weeks after treatment (**Fig 1A**), increase in pulmonary vascular resistance (**Fig 1B**), and increase in RV mass relative to the heart (**Fig 1C**), all consistent with a hemodynamically significant increase in pulmonary vascular resistance. In structural analysis, mice treated with L-NAME and HFD showed evidence of increased muscularization of small pulmonary arteries (< 50 μm) (**Fig 1D**) as well as increased medial wall thickness of vessels (**Fig 1E**).

**Figure 1.**
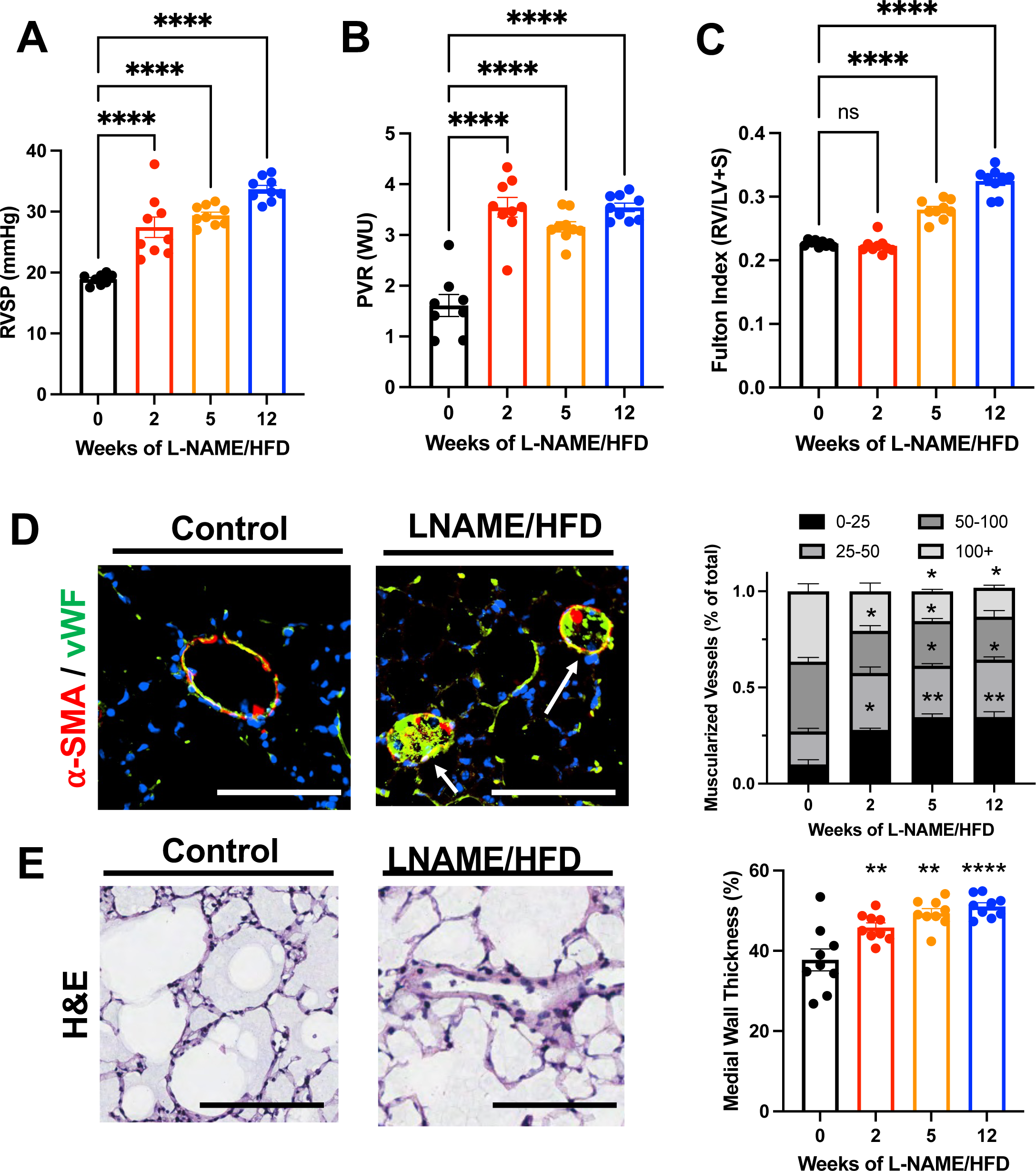
L-NAME and high fat diet treatment in mice causes (A) increased right ventricular systolic pressure, (B) increased pulmonary vascular resistance, (C) increased right ventricular mass relative to left ventricular and septal mass, (D) muscularization of small pulmonary vessels (< 50 μm) in lungs, and (E) increased medial wall thickness of pulmonary vessels in lungs. *\* p < 0.05, ** p < 0.01, *** p < 0.001, **** p < 0.0001 compared to baseline or control*.

### L-NAME/HFD Treated Mice Show Evidence of RV Dysfunction

Despite a normal stroke volume and cardiac output (**Fig 1A, Suppl Fig 1**), mice treated with L-NAME/HFD demonstrated an increase in the relaxation constant τ for the right ventricular (**Fig 2B**), decrease in end-systolic to end-arterial elastance ratio (Ees/Ea) (**Fig 2C**), increase in liver weight as a marker of right ventricular congestion (**Fig 2D**), and an increase in right ventricular end-diastolic pressure (**Fig 2E**). These findings were all suggestive of a load-stressed right ventricular with features of decompensation.

**Figure 2.**
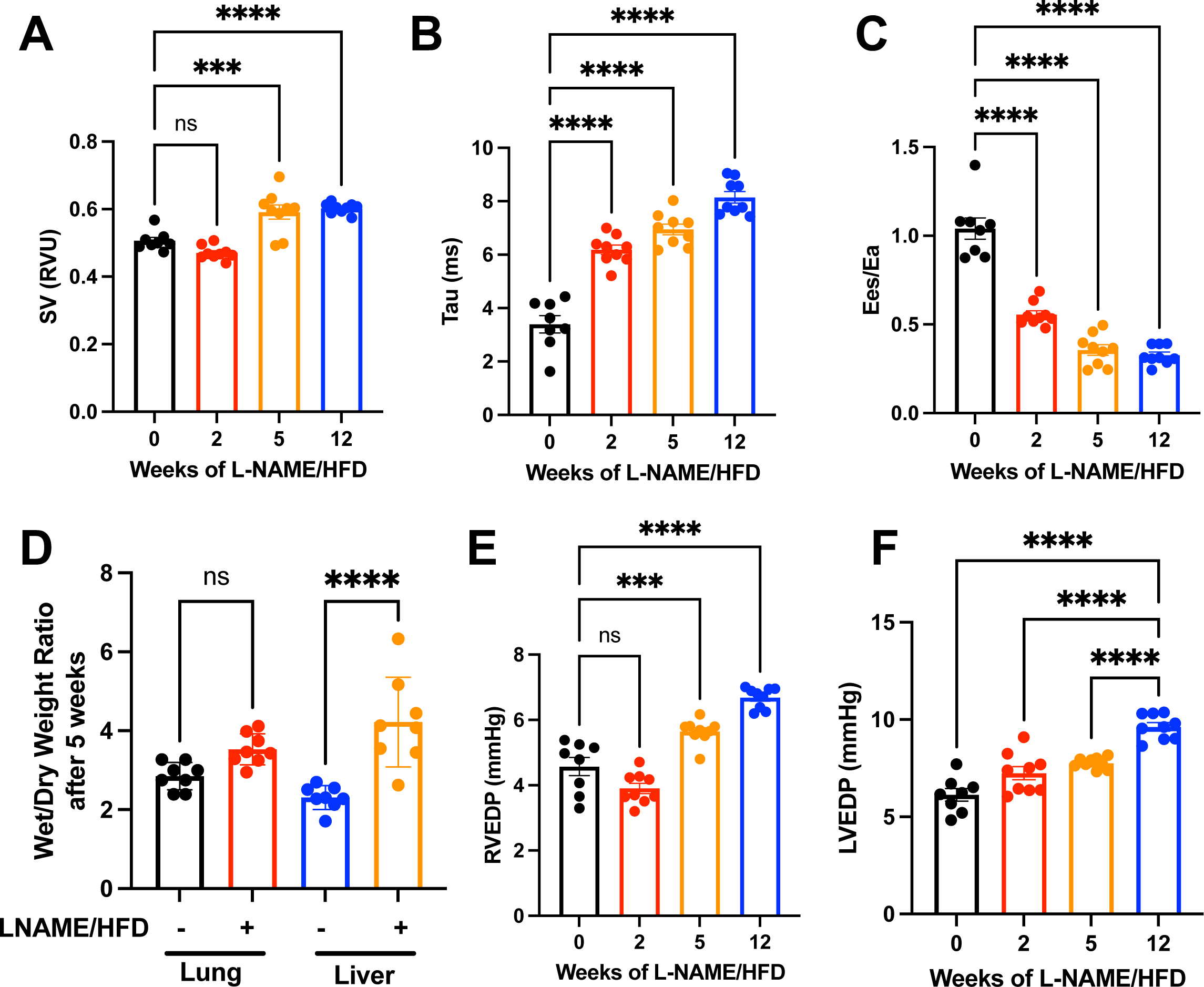
L-NAME and high fat diet treatment in mice (A) preserves stroke volume, (B) increased diastolic stiffness of the right ventricle as measured by the relaxation constant (τ), (C) decreased right ventricle-to-pulmonary artery coupling, (D) increased right heart congestion as measured by wet-to-dry liver mass, (E) increased right ventricular end-diastolic pressure, and (F) increased left ventricular end-diastolic pressure. *\* p < 0.05, ** p < 0.01, *** p < 0.001, **** p < 0.0001 compared to baseline or control*.

### L-NAME/HFD Treatment Results in Increased CD68+ Cells in Lungs

We next sought to better understand pathways that might contribute to pulmonary vascular remodeling after treatment with L-NAME and high fat diet. In mice treated with either L-NAME/HFD or control water/diet, RNA was isolated at 5 weeks after treatment and bulk RNA sequencing was performed to identify transcripts and gene ontologies that might be over-represented. In L-NAME/HFD treated lungs, the top 15 over-represented gene ontologies all represented pathways related to inflammation or immunity (**Fig 3A, Table S1**). Supporting these findings, many of the top transcripts that were differentially expressed included transcripts for cytokines such as Ccr7, Ccl5, Il12a, and Cxcl14 (**Fig 3B**), all known chemokines that affect macrophage polarization and recruitment. To better understand cell types that might be contributing to these changes, we next probed by RNA sequencing for transcript level expression of putative markers for T cells, B cells, monocyte/macrophages, and dendritic cells. Only CD68, a marker of macrophages, was found to be significantly increased (**Fig 3C**), while a trend towards increased expression was observed for other cell markers. We confirmed increased CD68 expression in lungs by Western blot (**Suppl Fig 3**). We also found an increase in CD68+ cells in lungs of L-NAME/HFD treated mice (**Fig 3D**) as well as in a second animal model of HFpEF (db/db mice). We confirmed that the db/db mice, which spontaneously develop HFpEF, also develop pulmonary hypertension (**Suppl Fig 4**). Finally, we also found an increase in CD68+ cells in the lungs sections of patients with hemodynamically confirmed PH with HFpEF who clinically underwent autopsies (**Fig 3E, Suppl Fig 5**).

**Figure 3.**
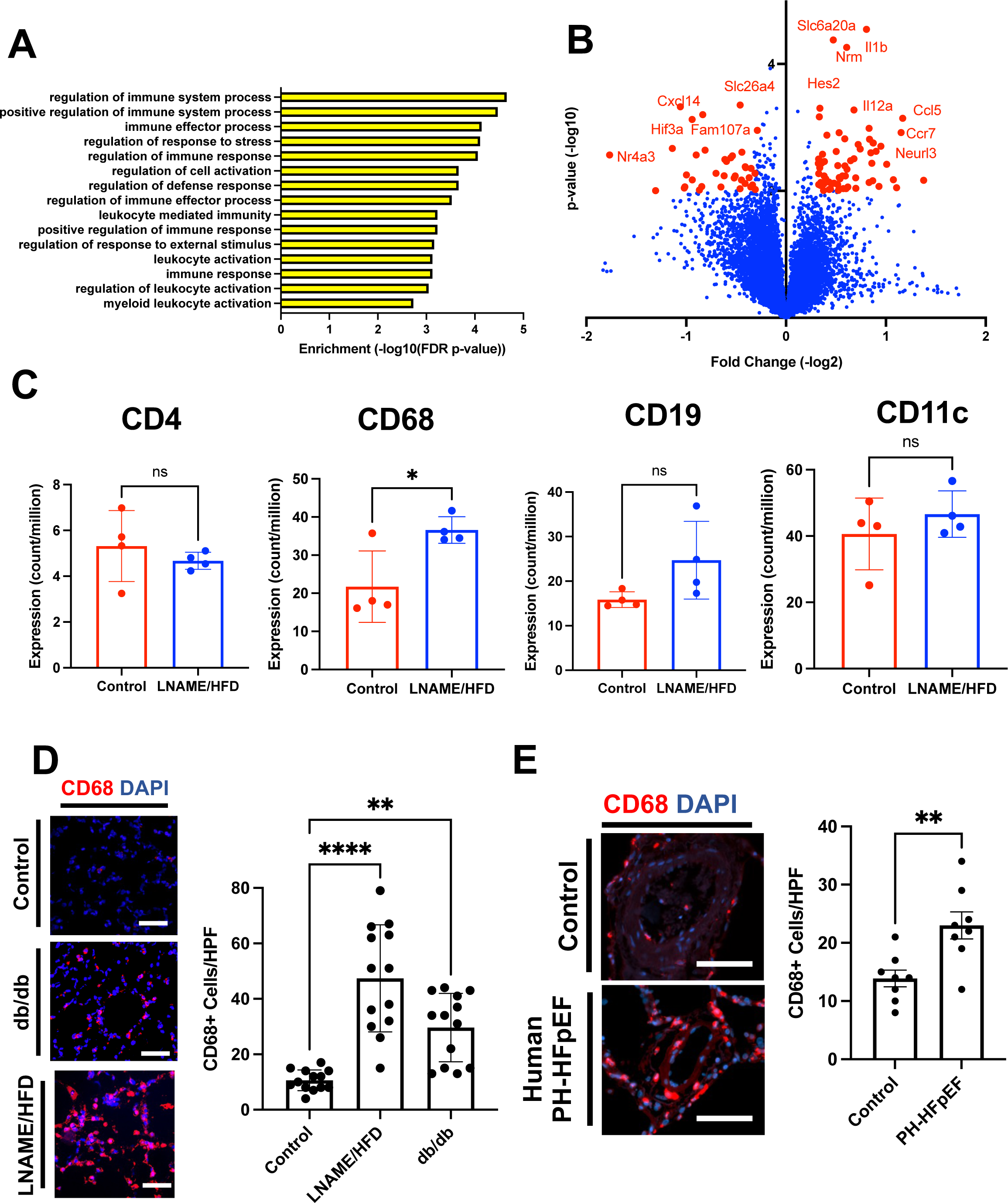
Bulk RNA sequencing of the whole lung in L-NAME and high fat diet (HFD) treated lungs compared to control identify that: (A) inflammatory gene ontologies are the top 15 most over-represented ontologies in the L-NAME/HFD treated lung, (B) transcript expression of cytokines represent the most differentially expressed transcripts, (C) transcript expression of macrophage marker CD68 is increased in the L-NAME/HFD treated lungs, (D) an increase in CD68+ cells in two animal models of heart failure with preserved ejection fraction, and (E) an increase in CD68+ cells in the lungs of patients with PH-HFpEF who underwent autopsy at death. *\* p < 0.05, ** p < 0.01, *** p < 0.001, **** p < 0.0001 compared to baseline or control*.

### L-NAME/HFD Treatment Results in Increased IL1β in Lungs

We next investigated whether L-NAME and high fat diet treatment affects the local and circulating cytokine proteome. By cytokine array, we found that cytokine expression differed in lung lysate compared to circulating plasma, suggesting that the inflammatory response observed in the lungs represented a local as opposed to systemic response (**Fig 4A, Suppl Fig 6**). The top 2 cytokines increased in the lung compared to plasma included CCL5 and IL1β (**Fig 4A**). To validate these exploratory findings, we confirmed an increase in protein expression of IL1β in the lungs of L-NAME/HFD treated mice by Western blot (**Fig 4B**) and ELISA (**Fig 4C**), but not CCL5 (**Suppl Fig 7**). Finally, we confirmed in collected plasma from patients undergoing clinically indicated right heart catheterization an increase in circulating IL1β levels in the venous blood, pulmonary arterial blood, and pulmonary capillary wedge blood of HFpEF patients with PH compared to non-PH controls (**Fig 4D**).

**Figure 4.**
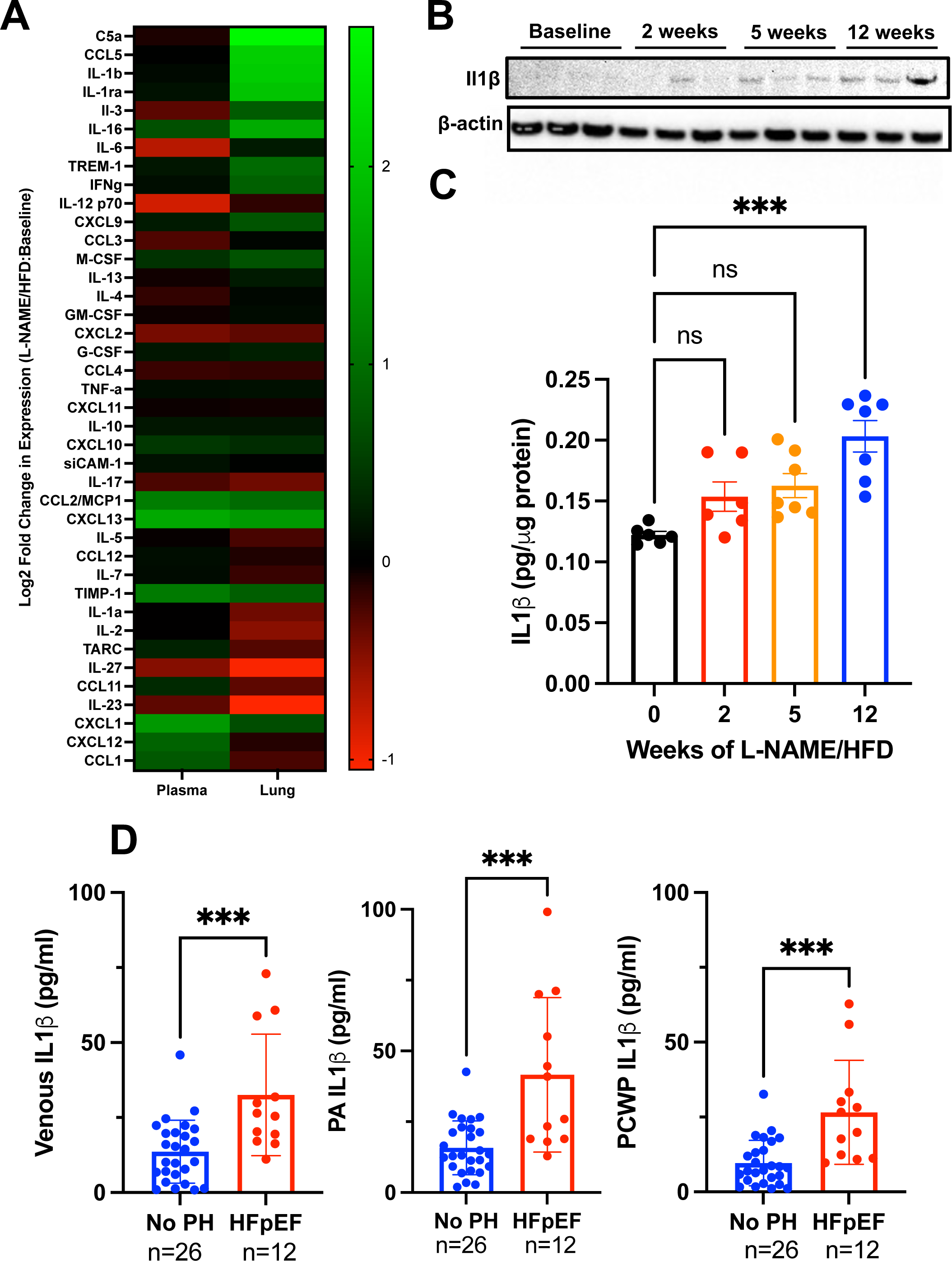
(A) Cytokine array in lungs and plasma of mice treated with L-NAME and high fat diet compared to control demonstrate lung-specific increases in cytokines and chemokines. (B) Western blot for IL1β confirms an increase in expression of mice treated with L-NAME and high fat diet. (C) ELISA for IL1β demonstrate an increase in expression in L-NAME/HFD treated mice. (D) IL1β levels by ELISA are increased in the venous, pulmonary artery, and pulmonary capillary wedge blood of patients with HFpEF compared to control patients without pulmonary hypertension. *\* p < 0.05, ** p < 0.01, *** p < 0.001, **** p < 0.0001 compared to baseline or control*.

### Monocyte/Macrophage Populations Display Pro-inflammatory Phenotype and Express IL1β

We next sought to better understand the phenotype and heterogeneity of the monocyte/macrophage populations in the lungs of L-NAME/HFD treated mice, and we additionally sought to better understand the potential sources of IL1β in the lung. To address these questions, we performed single cell RNA sequencing in lungs of mice treated with L-NAME/HFD or control water/diet as an efficient approach to answering these questions. By single cell RNA sequencing, we identified 30 cell populations within the lungs (**Fig 5A-B, Suppl Fig 8**). There were proportional increases in numbers of inflammatory cells in L-NAME/HFD treated mouse lungs including B-cell, T-cell, monocyte/macrophage, NK cell, and neutrophil populations (**Suppl Fig. 8**). Transcriptionally, monocyte, monocyte-dendritic cell, monocyte-derived macrophage, and activated monocyte populations showed an increase in transcript expression markers typically associated with M1-like macrophages compared to M2-like macrophages (**Fig 5C**), with similar transcript expression changes by bulk RNA sequencing in whole lung (**Suppl Fig 9**), whereas alveolar macrophage populations predominantly demonstrated a decrease in M1-associated transcripts and increase in M2-associated transcripts (**Fig 5C**). Nearly 90% of monocytes and 40% of activated monocyte, dendritic cells, and monocyte dendritic cells identified were Ccr2+, a marker of bone marrow derived monocyte/macrophage populations (**Fig. 5D**). Expression of IL1β transcripts was identified predominantly in neutrophil, activated monocyte, monocyte, and monocyte-derived macrophage populations (**Fig 5D**).

**Figure 5.**
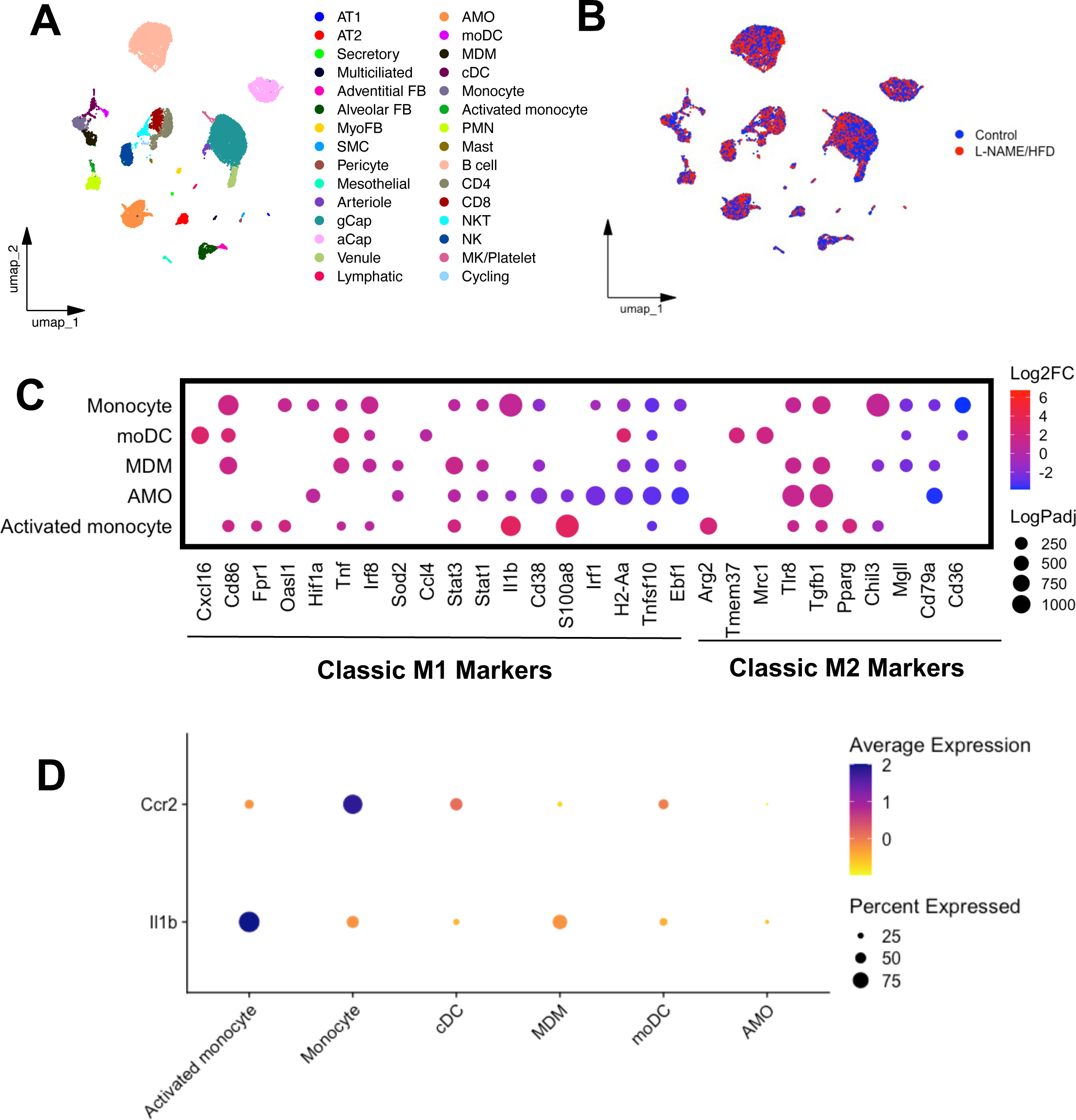
UMAP embedding of single cell RNA sequencing of lungs from mice treated with either L-NAME and high fat diet or control annotated by (A) cell type and (B) treatment group. (C) Transcript expression of classic M1 and M2 macrophage markers in various monocyte/macrophage populations in the lung. (D) Expression of Ccr2, a marker of bone marrow derived myeloid cells, and Il1β in various cell populations in the lung.

### Circulating Monocyte/Macrophages Contribute to Pulmonary Vascular Remodeling in HFpEF

Given the identification of increased bone marrow derived monocyte/macrophage populations in the lung, their pro-inflammatory transcriptome including IL1β expression in our PH-HFpEF model, we next sought to more directly test the hypothesis that circulating monocyte/macrophage populations contribute to pulmonary vascular remodeling in the L-NAME/HFD model of HFpEF with PH. To deplete the circulating source of monocytes and macrophages, we treated mice with L-NAME/HFD for 5 weeks and administered twice weekly injections of either clodronate or PBS containing liposomes (**Fig 6A**). Clodronate treatment resulted in reduced CD68 expression in mouse lungs (**Fig 6B)** as well as lower IL1β levels by ELISA (**Fig 6C**). Mice treated with clodronate compared to PBS liposomes also demonstrated less weight gain, near normal RV systolic pressure, normal RV mass compared to LV and septal mass in the heart, near normal RV end-diastolic pressure, and a decrease in muscularization of small vessels histologically in the lungs (**Fig 6D,E**), all suggestive that a bone marrow derived source of monocytes/macrophages contributes to pathologic pulmonary vascular remodeling and pulmonary hypertension in the L-NAME/HFD model of HFpEF.

**Figure 6.**
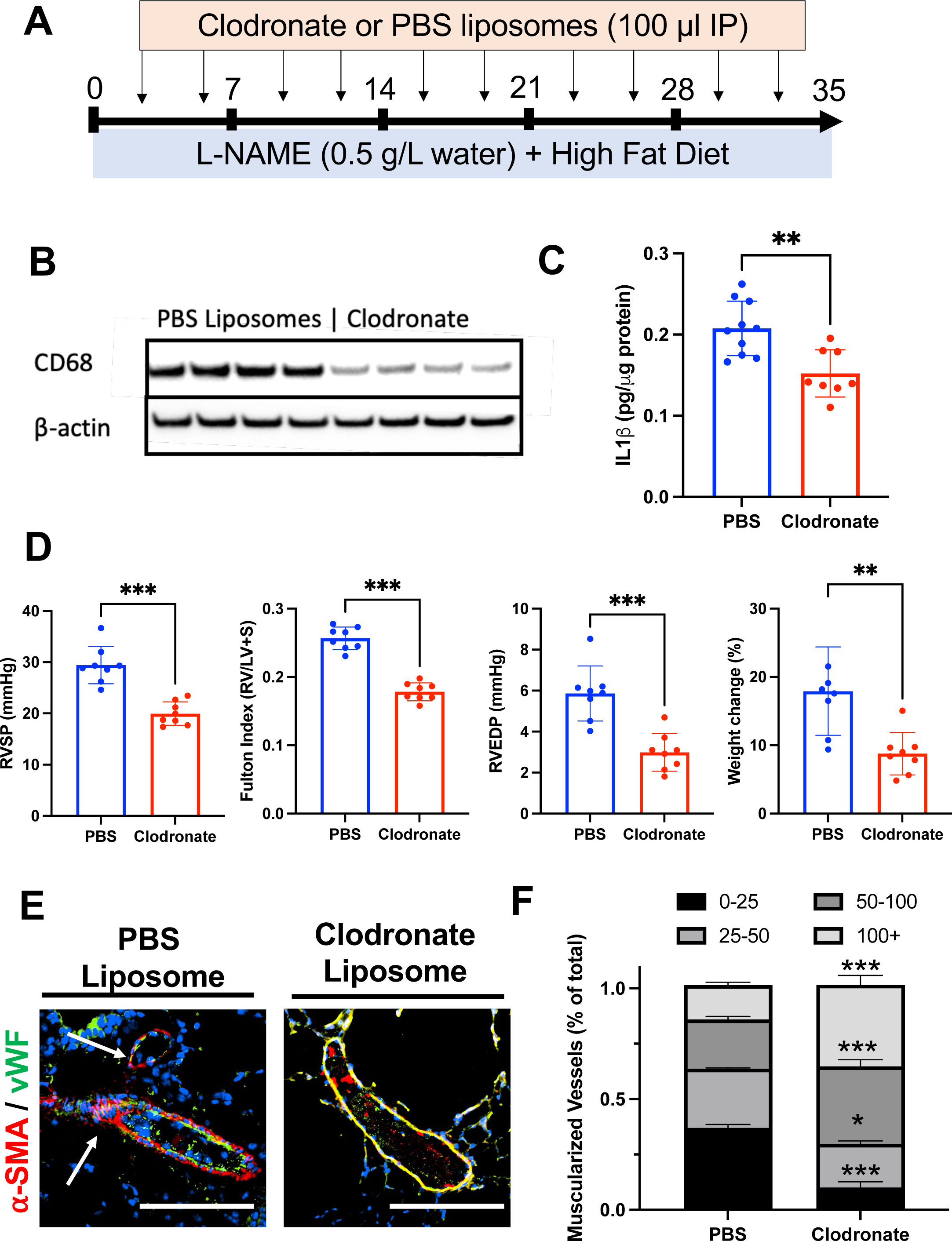
(A) Schematic depiction of clodronate-mediated monocyte/macrophage depletion studies. (B) CD68 expression by Western blot in the lung after clodronate or PBS liposome treatment. (C) IL1β levels by ELISA in the whole lung after clodronate or PBS liposome treatment. (D) Right ventricular systolic pressure, right ventricular mass relative to left ventricular and septal mass, right ventricular end-diastolic pressure, and weight change in mice after clodronate vs PBS liposome treatment. (E) Muscularization of small vessels in the lungs decrease after clodronate treatment in L-NAME and high fat diet treated mice. *\* p < 0.05, ** p < 0.01, *** p < 0.001, **** p < 0.0001 compared to baseline or control*.

### IL1β Contributes to Pulmonary Vascular Remodeling in HFpEF

We finally sought to test the hypothesis that IL1β contributes to pulmonary vascular remodeling in response to L-NAME/HFD treatment. To mimic a treatment or reversal model that would be more clinically relevant, we first treated mice with L-NAME and HFD for 3 weeks, followed by continuation of L-NAME/HFD with concomitant administration of either IL1β neutralizing antibody or isotype control (**Fig 7A**). Treatment with IL1β neutralizing antibody resulted in a decrease in IL1β levels by ELISA in the lung (**Fig 7B**), decreased RV systolic pressure (**Fig 7C**), and decreased RV mass as compared to LV and septal mass (**Fig 7D**). No changes in RV end-diastolic pressure or weight gain were observed (**Fig 7E,F**). Histologically, IL1β depletion also resulted in a decrease in small vessel muscularization (**Fig 7G**). Based on our previous data supporting myeloid cells as the primary source of IL1β, our findings suggest that myeloid-derived IL1β contributes to PH-HFpEF.

**Figure 7.**
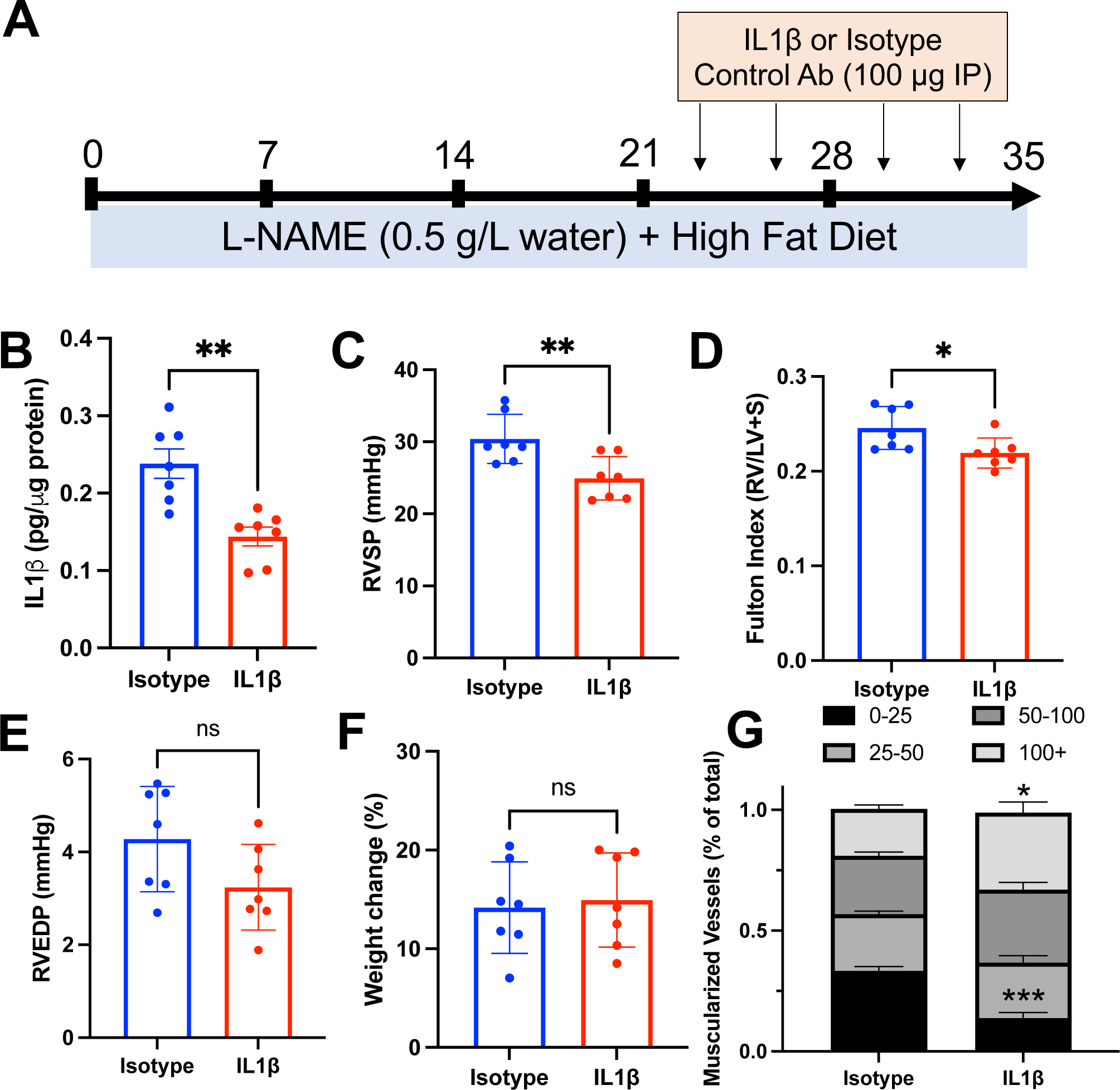
(A) Schematic depiction of IL1β antibody mediated depletion studies. (B) Il1β levels by ELISA, (C) right ventricular systolic pressure, (C) right ventricular end-diastolic pressure, weight change, and (E) muscularization of vessels in the lung after treatment with IL1β specific or isotype control antibody in mice treated with L-NAME and high fat diet. *\* p < 0.05, ** p < 0.01, *** p < 0.001, **** p < 0.0001 compared to baseline or control*.

## DISCUSSION

Pulmonary vascular remodeling is a highly morbid condition that commonly occurs in HFpEF, but pathways that drive remodeling in HFpEF are poorly understood. The present study demonstrates that a widely accepted multi-hit model of metabolically driven HFpEF, induced by the combination of L-NAME and high fat diet (HFD) administration, also shows features of pulmonary vascular remodeling and pulmonary hypertension that are often found in patients with HFpEF.^2^ Through the use of discovery based approaches and validation from both a second animal model of HFpEF and human lung and plasma samples, we also show that circulating monocyte/macrophage populations and IL1β contribute to pathologic pulmonary vascular remodeling in HFpEF.

Heart failure is estimated to affect nearly 10% of the US population throughout their lifetime, approximately half of which constitutes HFpEF and carries a 75% 5-year mortality despite no current available therapies that improve mortality in this population.^14, 15^ The coexisting presence of pulmonary hypertension, which variably occurs in up to 80% of patients with HFpEF,^16^ is independently associated with worse prognosis.^17^ To date, however, strategies to prevent or reverse PH in HFpEF have been hampered by a limited understanding of the pathogenesis underlying pathologic pulmonary vascular remodeling in HFpEF, which has been identified as a key research priority for the study of HFpEF.^3, 18^

Our study adds to a growing literature of available models that approximate features of pathologic pulmonary vascular remodeling in HFpEF.^19–21^ The L-NAME/HFD model has the distinct advantages that it is technically straightforward to implement, it replicates known deficits in nitric oxide and PKG signaling from human studies of HFpEF,^22, 23^ recapitulates many other extra-cardiac features of HFpEF including metabolic disease that is prominent in the human condition,^24^ and is very amenable to further mechanistic studies since it is implemented in a mouse organism for which many transgenic models already exist.

Discovery based approaches in our study found inflammatory pathways to be altered in the remodeling lung of PH-HFpEF mice. Specifically, we focused on the role of recruited monocyte/macrophage populations and IL1β in promoting pathologic pulmonary vascular remodeling. Beyond corroboration in both human samples as well as autopsy-derived lung slides from patients with PH-HFpEF, our findings also mirror the findings from both human studies and pre-clinical animal studies that have also implicated inflammatory pathways in the pathogenesis of PH in HFpEF including IL1β.^19, 25^ The role of IL1β in regulating endothelial cell permeability and lung injury is well established,^26, 27^ and activation of the NLRP3 inflammasome (upstream of IL1β expression and release) has been shown to regulate PH and RV failure in other forms of PH not related to heart failure as well.^28^ Finally, studies have also found that macrophage-mediated IL1β partially regulates cardiac remodeling in the animal models of HFpEF.^29^ Our study expands upon these initial findings by demonstrating that the extracardiac manifestation of pulmonary vascular remodeling is also partially regulated by monocyte/macrophage populations and IL1β. Our study showed that the cytokine profile of the lung differed from that of the plasma in our animal model of HFpEF, however. These findings suggest that organ-specific inflammation in HFpEF may be modified, in part, by the microenvironment within specific organs, however further work is necessary to directly test this hypothesis.

There are limitations to our work. While our study focused on the role of monocyte/macrophages and IL1β in regulating pulmonary vascular remodeling in the L-NAME/HFD model of HFpEF, our findings also identify other inflammatory cell types and cytokines that are present within the lung. We cannot exclude the contribution of other cell types and cytokines to pulmonary vascular remodeling. Furthermore, while our plasma cytokine profile significantly differed from the lung cytokine profile, we cannot definitively exclude the possibility that extrapulmonary effects of monocyte/macrophage or IL1β depletion regulate pulmonary vascular remodeling through interorgan communication. Finally, our studies demonstrated that other cell sources such as neutrophils also expressed IL1β and likely contributed to IL1β-mediated pulmonary vascular remodeling. While studies have shown that clodronate-mediated depletion of monocyte/macrophage populations in the lung can also reduce neutrophil recruitment and activation,^30^ we cannot definitively rule out the contribution of neutrophil derived IL1β to pulmonary vascular remodeling in HFpEF.

In summary, our study demonstrates that the L-NAME/HFD model of HFpEF also recapitulates features of pulmonary vascular remodeling that are observed in HFpEF patients, and we further demonstrate the role of monocyte/macrophage populations and IL1β in regulating pulmonary vascular remodeling in HFpEF. These findings not only provide novel insight into the elusive pathophysiology of pulmonary vascular remodeling in heart failure, but also raise important questions regarding the interplay between metabolic syndrome-driven cardiac disease and pulmonary vascular inflammation. Particularly with the growth of anti-inflammatory therapies in the treatment of cardiovascular disorders such as IL-1β targeting therapies, and recent insights into the immunomodulatory properties of more contemporary anti-diabetes therapies such as GLP-1 agonists and SGLT2 inhibitors, the PH-HFpEF model induced by L-NAME+HFD diet treatment in mice may serve as an important vehicle for not only identifying relevant pathophysiologic mechanisms in PH-HFpEF but also testing promising potential therapies.

---

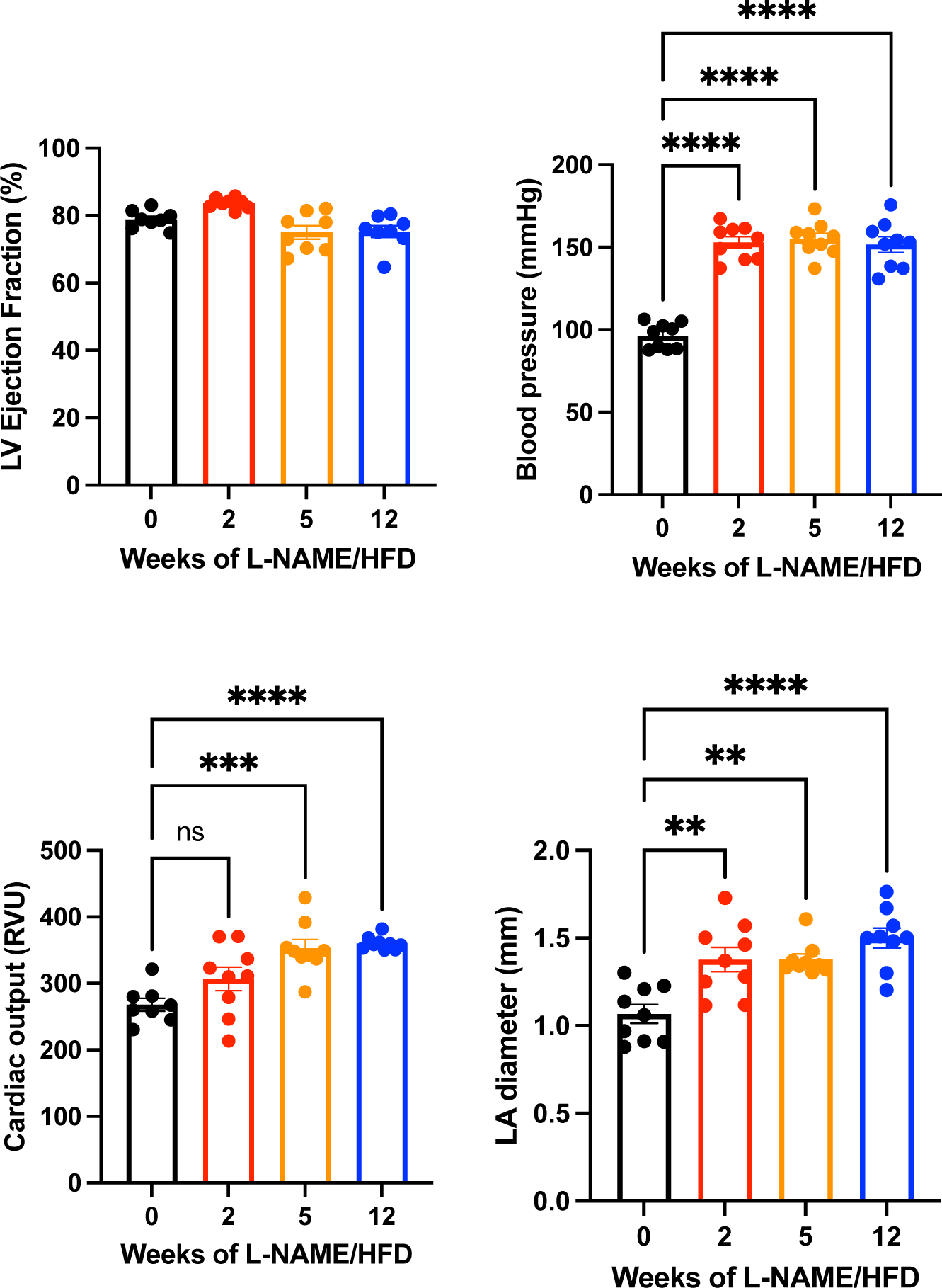
Supplementary Figure 1.

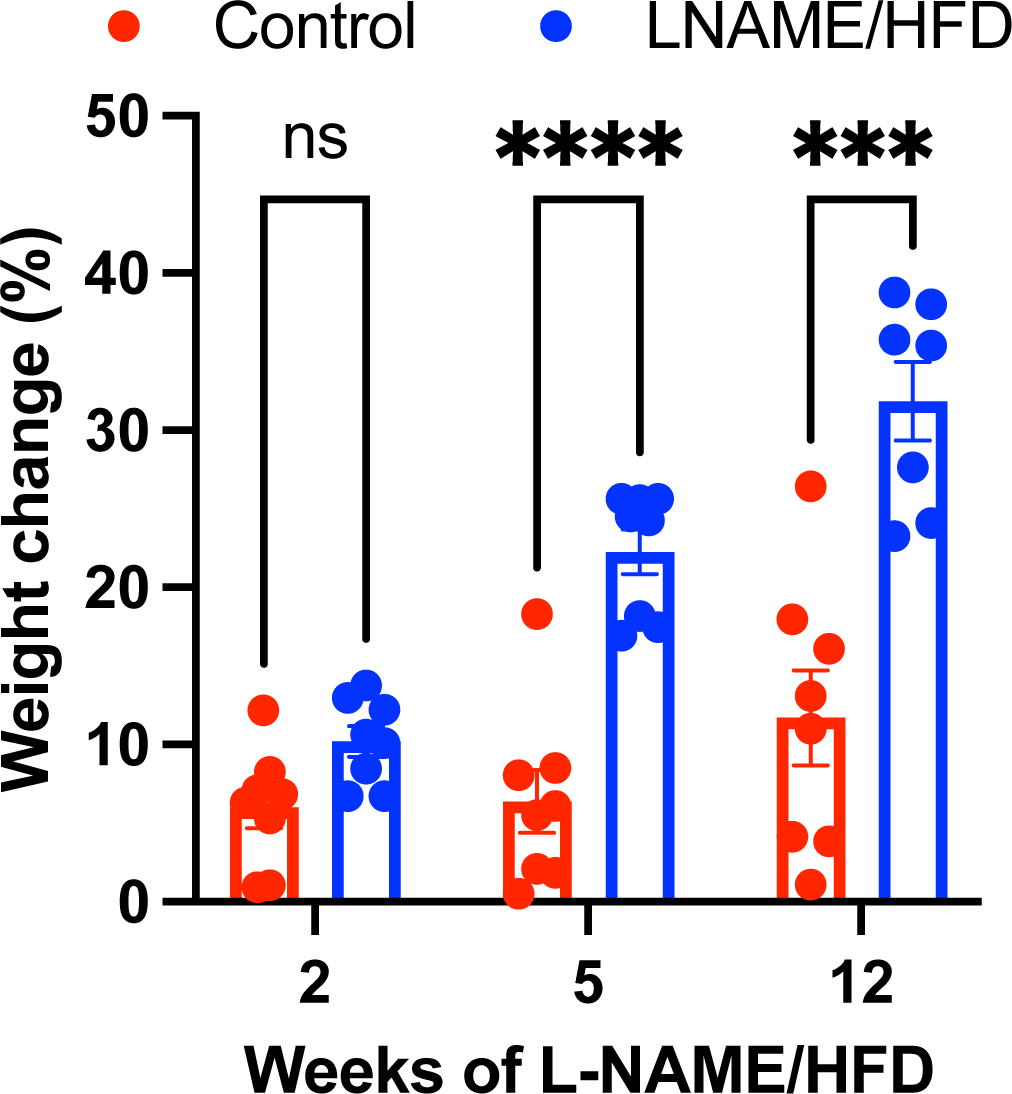
Supplementary Figure 2.

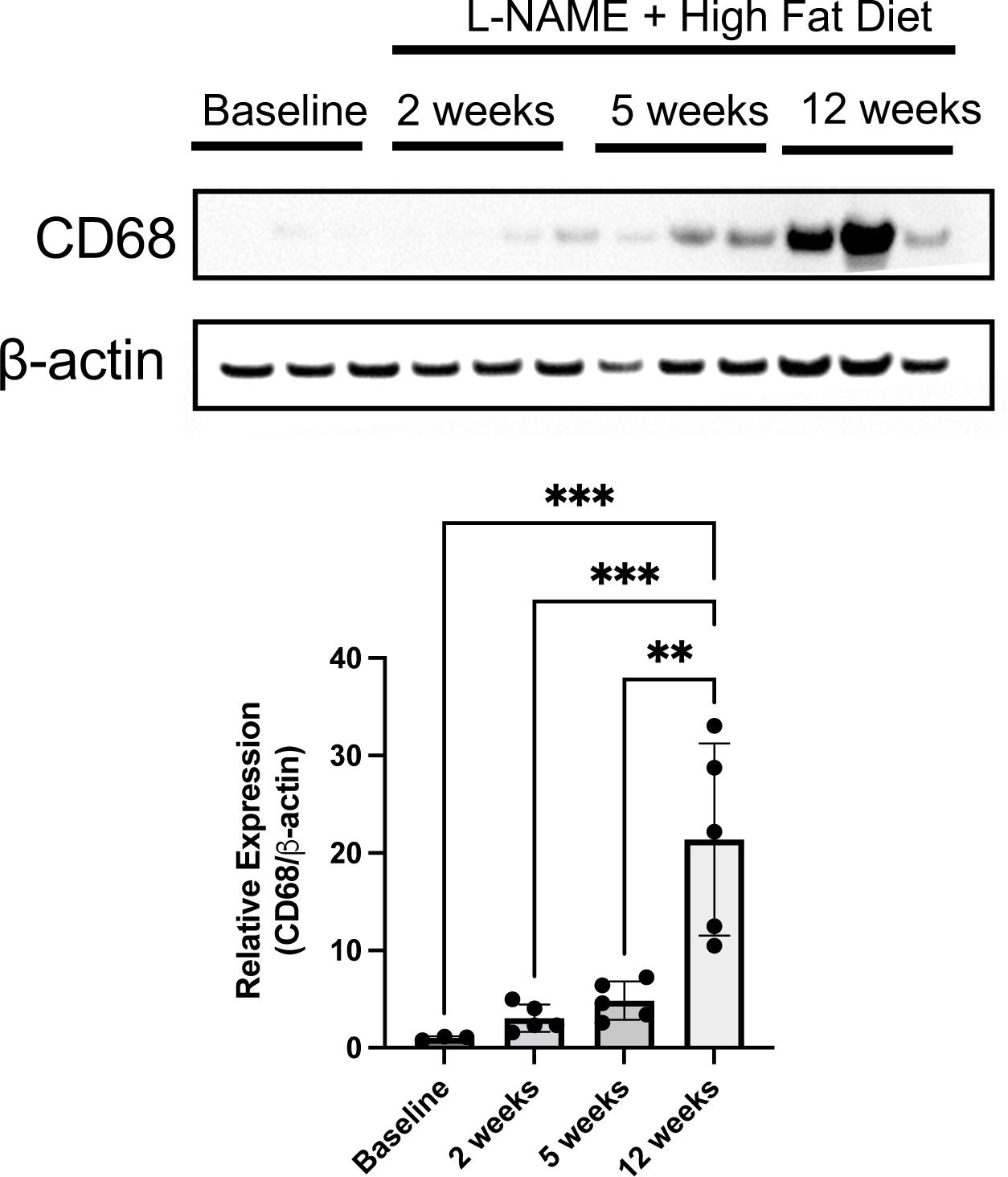
Supplementary Figure 3.

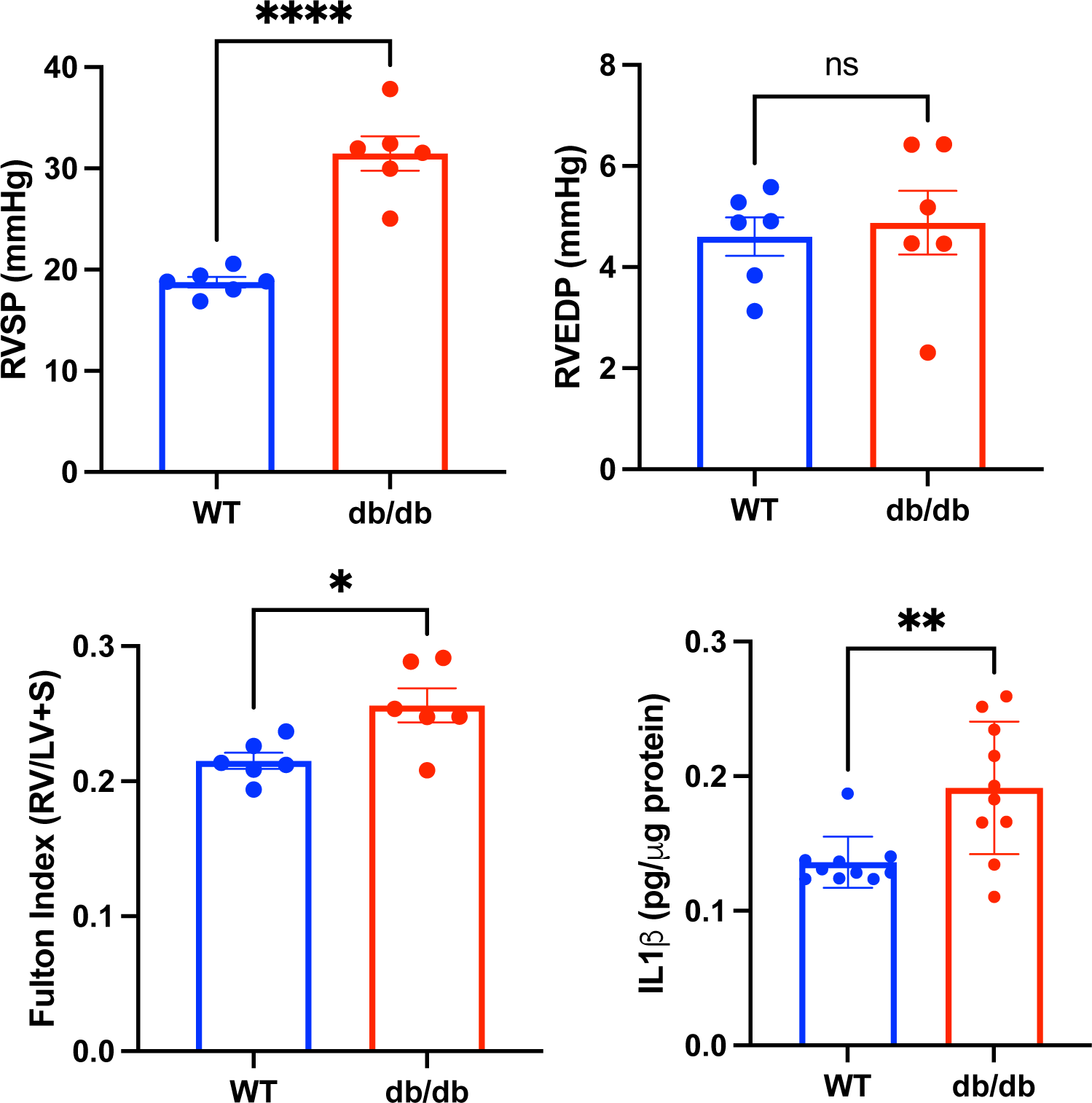
Supplementary Figure 4.

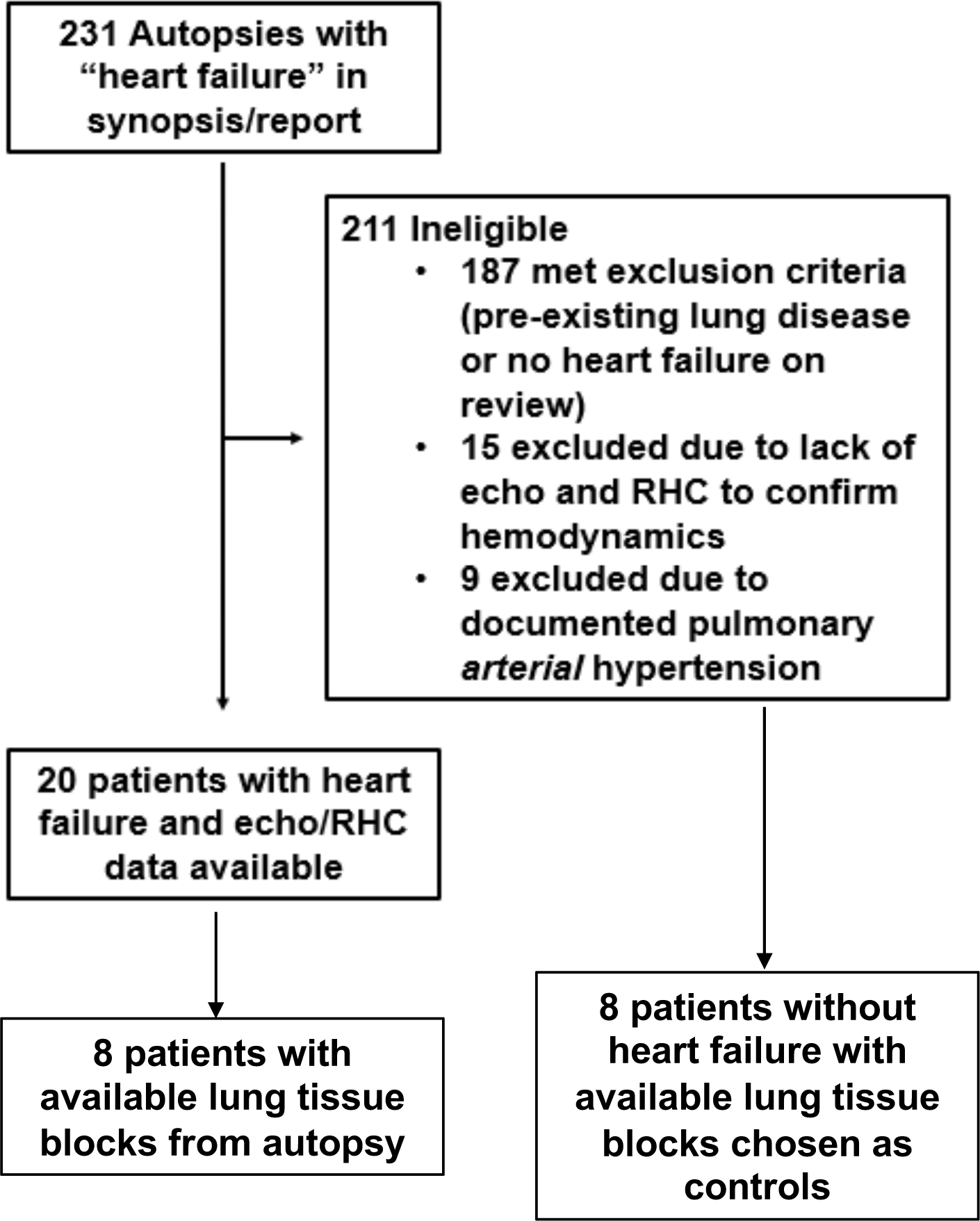
Supplementary Figure 5.

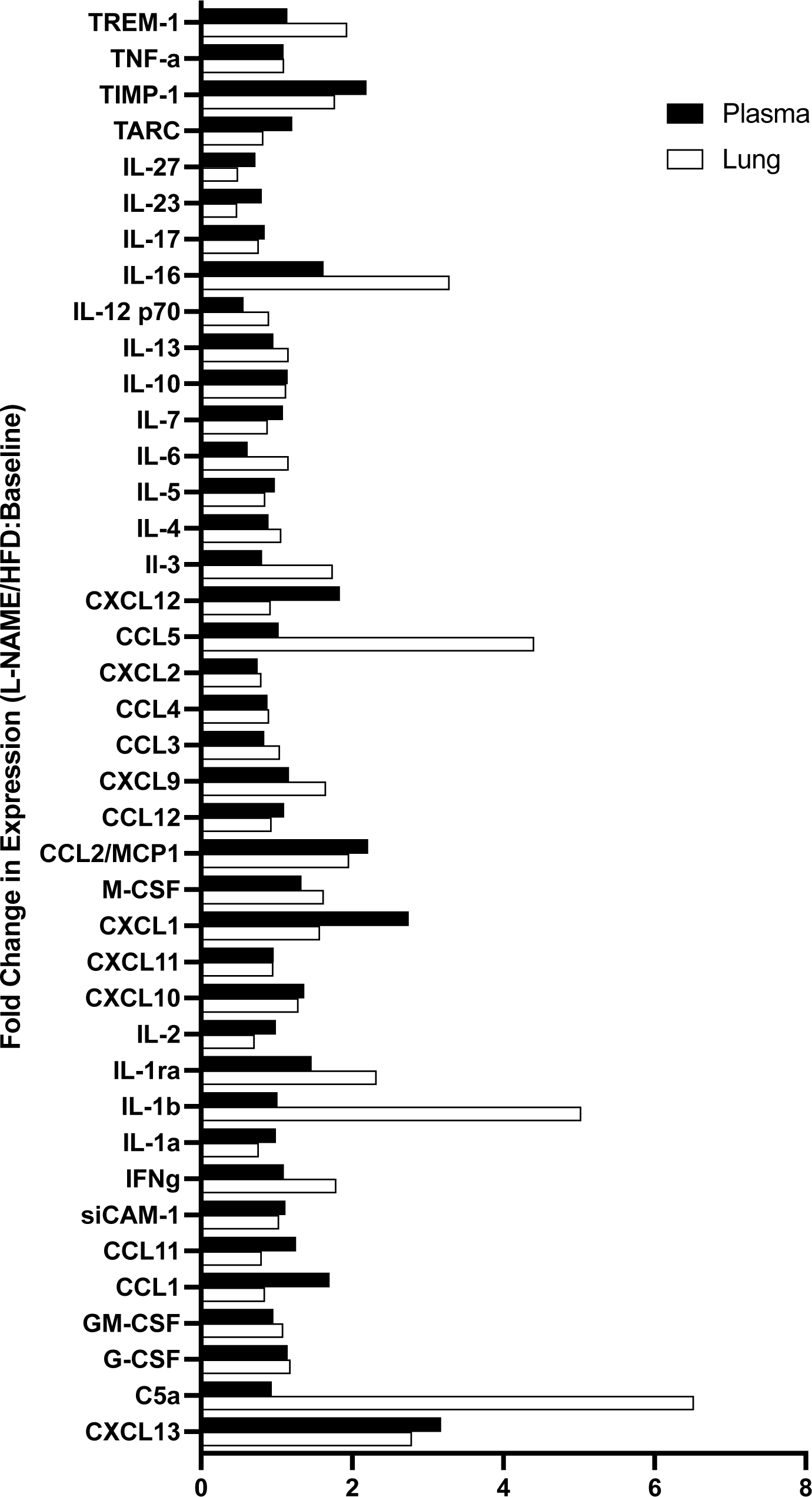
Supplementary Figure 6.

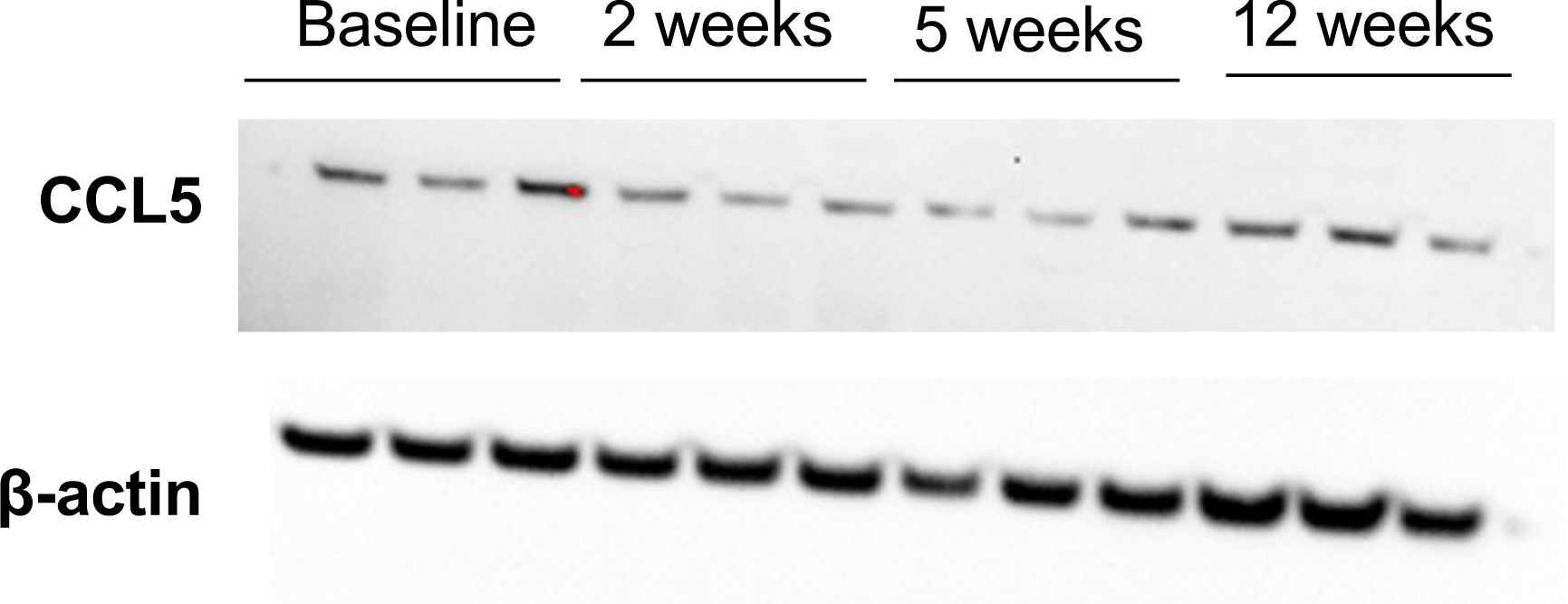
Supplementary Figure 7.

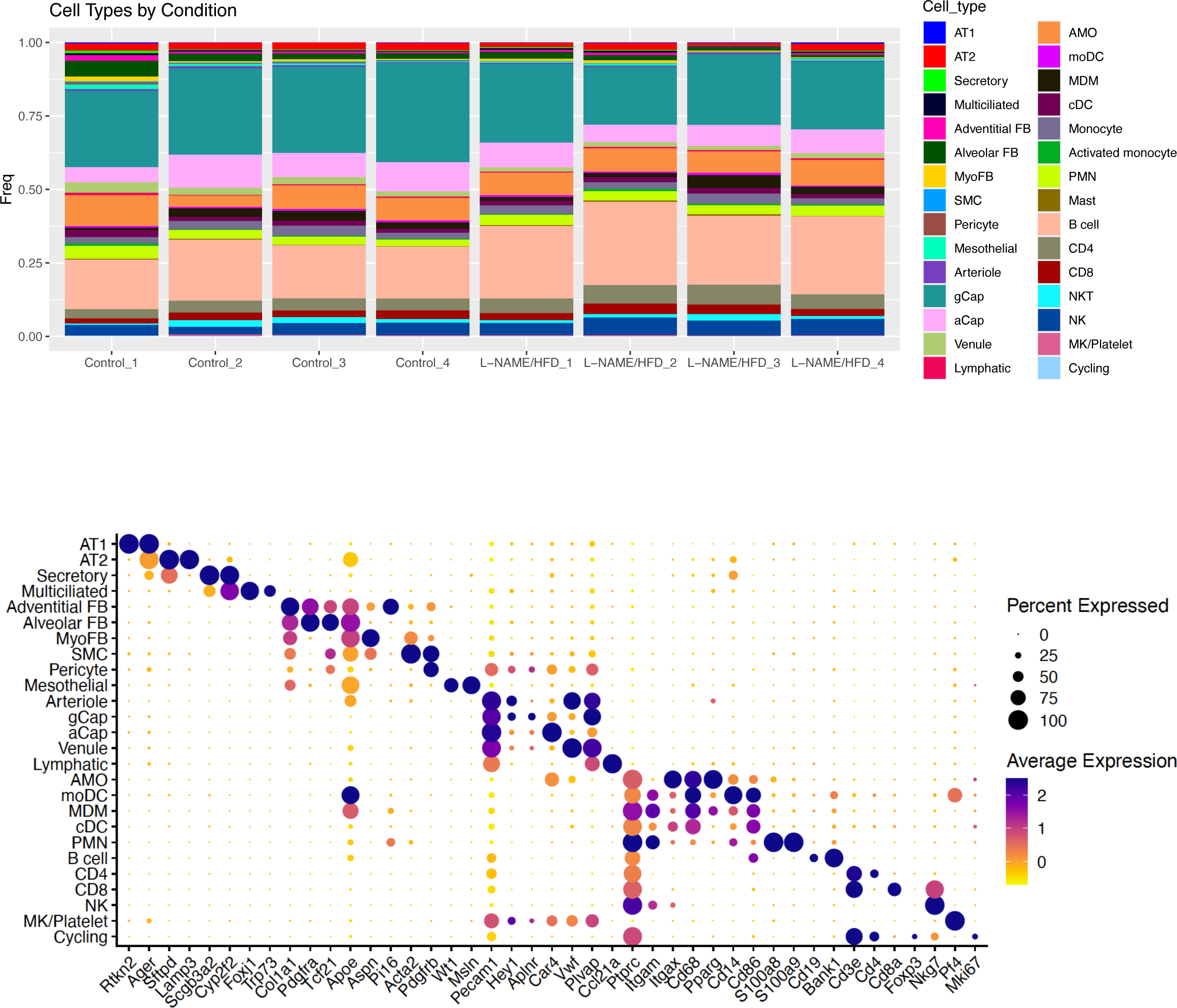
Supplementary Figure 8.

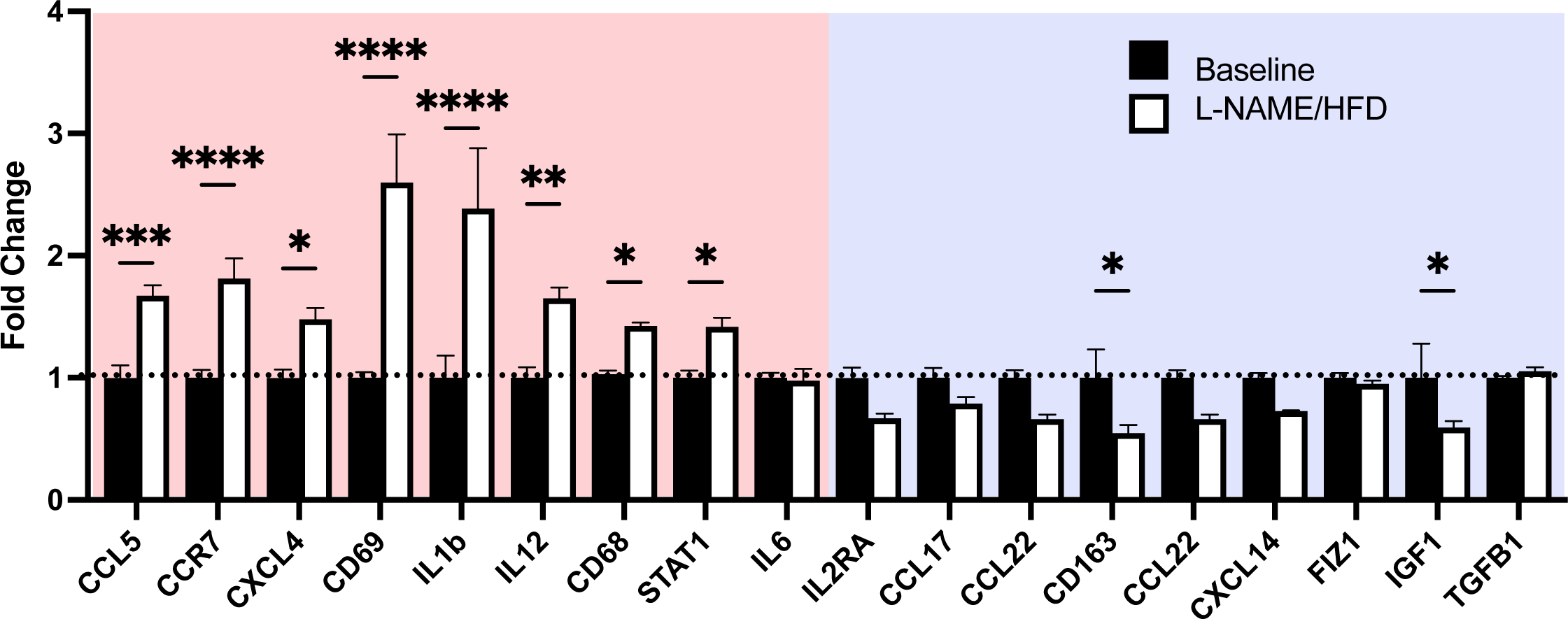
Supplementary Figure 9.

**Table S1.** Over-represented gene ontologies in lungs of mice treated with L-NAME and high fat diet compared to control water and diet at 5 weeks.

|  |  |  |  |  |  |  |  |
| --- | --- | --- | --- | --- | --- | --- | --- |
| GO | Description | Size of Ontology | Overlapping Genes | Expected Overlap | Enrichment Ratio | P-value | FDR p-value |
| GO:0002682 | regulation of immune system process | 1230 | 32 | 9.853483715 | 3.247582371 | 2.45E-09 | 2.20E-05 |
| GO:0002684 | positive regulation of immune system process | 879 | 26 | 7.041635923 | 3.692323813 | 7.45E-09 | 3.36E-05 |
| GO:0002252 | immune effector process | 678 | 22 | 5.431432487 | 4.050496817 | 2.41E-08 | 7.24E-05 |
| GO:0080134 | regulation of response to stress | 1227 | 30 | 9.829450827 | 3.052052503 | 3.45E-08 | 7.78E-05 |
| GO:0050776 | regulation of immune response | 705 | 22 | 5.647728471 | 3.895371407 | 4.80E-08 | 8.66E-05 |
| GO:0001775 | cell activation | 893 | 24 | 7.153789396 | 3.354863327 | 1.81E-07 | 2.16E-04 |
| GO:0050865 | regulation of cell activation | 521 | 18 | 4.173711395 | 4.312708354 | 2.01E-07 | 2.16E-04 |
| GO:0031347 | regulation of defense response | 580 | 19 | 4.646358174 | 4.089224138 | 2.05E-07 | 2.16E-04 |
| GO:0042592 | homeostatic process | 1809 | 36 | 14.49183093 | 2.484158157 | 2.15E-07 | 2.16E-04 |
| GO:0002697 | regulation of immune effector process | 373 | 15 | 2.988088964 | 5.019930859 | 3.33E-07 | 3.00E-04 |
| GO:0002443 | leukocyte mediated immunity | 345 | 14 | 2.763782017 | 5.065522502 | 7.52E-07 | 5.90E-04 |
| GO:0050778 | positive regulation of immune response | 572 | 18 | 4.582270475 | 3.928183658 | 7.85E-07 | 5.90E-04 |
| GO:0032101 | regulation of response to external stimulus | 708 | 20 | 5.671761358 | 3.526241451 | 9.92E-07 | 6.88E-04 |
| GO:0045321 | leukocyte activation | 784 | 21 | 6.280594498 | 3.343632519 | 1.22E-06 | 7.49E-04 |
| GO:0006955 | immune response | 1293 | 28 | 10.35817434 | 2.703179061 | 1.25E-06 | 7.49E-04 |
| GO:0022407 | regulation of cell-cell adhesion | 369 | 14 | 2.956045114 | 4.736057624 | 1.66E-06 | 8.94E-04 |
| GO:0002694 | regulation of leukocyte activation | 482 | 16 | 3.861283862 | 4.143699498 | 1.69E-06 | 8.94E-04 |
| GO:0002274 | myeloid leukocyte activation | 194 | 10 | 1.5541267 | 6.434481823 | 3.78E-06 | 0.001848402 |
| GO:0002250 | adaptive immune response | 397 | 14 | 3.180352061 | 4.402028371 | 3.89E-06 | 0.001848402 |
| GO:0051707 | response to other organism | 942 | 22 | 7.546326552 | 2.915325735 | 6.17E-06 | 0.002693107 |
| GO:0043207 | response to external biotic stimulus | 945 | 22 | 7.570359439 | 2.906070732 | 6.49E-06 | 0.002693107 |
| GO:0032102 | negative regulation of response to external stimulus | 305 | 12 | 2.443343523 | 4.911302847 | 6.57E-06 | 0.002693107 |
| GO:0002764 | immune response-regulating signaling pathway | 310 | 12 | 2.483398335 | 4.832088285 | 7.75E-06 | 0.003036618 |
| GO:0002449 | lymphocyte mediated immunity | 261 | 11 | 2.090861178 | 5.260990119 | 8.47E-06 | 0.003099585 |
| GO:0002253 | activation of immune response | 368 | 13 | 2.948034152 | 4.409718249 | 8.59E-06 | 0.003099585 |
| GO:0045088 | regulation of innate immune response | 267 | 11 | 2.138926953 | 5.142765622 | 1.05E-05 | 0.003440007 |
| GO:0072676 | lymphocyte migration | 94 | 7 | 0.753030463 | 9.295772676 | 1.05E-05 | 0.003440007 |
| GO:0009607 | response to biotic stimulus | 977 | 22 | 7.826710235 | 2.810887249 | 1.09E-05 | 0.003440007 |
| GO:0012501 | programmed cell death | 1874 | 33 | 15.01254348 | 2.198161827 | 1.11E-05 | 0.003440007 |
| GO:0030155 | regulation of cell adhesion | 639 | 17 | 5.119004954 | 3.320957911 | 1.49E-05 | 0.004488588 |

## Supplemental Methods

### Mouse Echocardiography and Catheterization

Mouse echocardiography and catheterization was performed as previously reported.^1^ One day prior to catheterization, mice were anesthetized with 2-3% isoflurane. Depilatory cream was applied to the thorax. Mice were then placed on a heated table in supine position and images were acquired using the VisualSonics Vevo700 Platform and 707B transducer (30 MHz) in B-mode, M-mode, and Doppler mode. Parasternal long-axis, short-axis, and modified RV-centric views were obtained as previously described.^2^ Thereafter, mice were allowed to recover for 24 hours before undergoing anesthesia again with 2-3% isoflurane for open chest catheterization. Mice were orotracheally intubated with 22g catheter, mechanically ventilated at 18 cc/kg, and anesthetized with 2-3% vaporized isoflurane general anesthesia. On a heated surgical table in supine, ventral side-up position, a vertical incision was made along the linea alba in the rectus abdominus sheath with scissors and cautery, and extended laterally. Cautery was then used to take down the diaphragm and exposure the thorax. A 1.4 French Mikro-tip catheter was directly inserted into the left ventricle for measurement of pressure and volume within the left ventricle. Afterwards, the catheter was removed and directly inserted into the right ventricle. Hemostasis of the left ventricle was ensured prior to catheterization of the right ventricle to ensure no effect of volume loss on findings. Hemodynamics were continuously recorded with a Millar MPVS-300 unit coupled to a Powerlab 8-SP analog-to-digital converter acquired at 1000 Hz and captured to a Macintosh G4 (Millar Instruments, Houston, TX). Thereafter, direct cardiac puncture was used to phlebotomize mice using a heparinized syringe and mice were sacrificed with cervical dislocation. Tissue and samples were then harvested for downstream studies from mice, including morphometric measurements. Volume was measured in relative volume units based on recommended calibrations from Millar Instruments. Pulmonary vascular resistance was calculated in Wood units using the transpulmonary gradient to cardiac output ratio, as measured by catheterization.

### Plasma Sample Collection from Patients

Samples were analyzed from a previous prospective, cross-sectional study.^3, 4^ The study was approved by the Vanderbilt University Institutional Review Board and all participants provided written informed consent. Inclusion criteria included patients aged 18 or older, referral for right heart catheterization with or without concomitant left heart catheterization as part of their usual care. Exclusion criteria included LVEF ≤ 40%, atrial fibrillation on the day of catheterization, significant anemia (hemoglobin < 10 g/dL and hematocrit < 30%), pregnancy, and treatment with PAH-specific medications or nitrates. Samples were collected after Swan-Ganz catheter placement in the proximal pulmonary artery or wedge position, confirmed by pressure waveform analysis. A 5-10 cc blood sample was collected with minimal aspiration pressure, immediately placed on ice, and then hand-delivered to the Vanderbilt Core Laboratory for Cardiovascular Translational and Clinical Research where plasma was separated, aliquoted, and stored in a −80 °C freezer for analysis. For classification of cases, all hemodynamic tracings were independently reviewed by cardiologists (DFM and KM), and the following definitions were used for classification: (1) PH-HFpEF: mPAP ≥ 20 mmHg and pulmonary artery wedge pressure > 15 mmHg) vs o(2) no PH (mPAP < 20 mmHg). All pressures were measured at end expiration at rest in supine patients. Biomarker analysis for IL1β was conducted following manufacturer’s recommended protocol on samples (Thermo, A35574, Waltham, MA) in triplicate on each sample.

### Lung Autopsy Slide Analysis

To identify clinically conducted autopsies in patients with pulmonary hypertension and heart failure, we interrogated Vanderbilt University Medical Center’s clinical autopsy database. Autopsy cases between 2013 and 2018 were initially screen for the presence of terms “heart failure” in the case synopsis or pathology report with additional filters of patient aged 18 or older. Cases were then individually reviewed (VA). Inclusion criteria included any pre-mortem right heart catheterization performed that demonstrated mean PA pressure > 20 mmHg and pulmonary arterial wedge pressure > 15 mmHg as well as echocardiography-based on cardiac MRI-based measurement of left ventricular ejection fraction ≥ 50%. Pre-determined exclusion criteria included: (1) known pre-existing lung disease such as obstructive lung disease as this would alter lung histology, (2) the presence of a primary pulmonary vascular disease such as pulmonary arterial hypertension or pulmonary veno-occlusive disease, (3) cause of death that was deemed to be due to acute lung injury (ARDS) or infection (pneumonia), (4) presence of congenital heart disease, and (4) presence of an autoimmune condition or immunosuppressive therapy for any indication as this may alter vascular remodeling via alteration of inflammatory pathways. A total of 231 autopsy cases were reviewed, of which 20 met inclusion criteria for PH and HFpEF without any exclusion criteria. Of those, 8 still had available blocks of lung tissue, which were used for the final analysis. An additional 8 lung tissue sections were randomly chosen among excluded autopsies of patients without heart failure.

### Western Blot and Immunostaining

Lung slides, both mouse and human, were deparaffinized and underwent heat antigen retrieval for 20 min in Tris EDTA pH 9 antigen retrieval buffer. Samples were blocked for 2 h in 10% goat serum and 1% bovine serum albumin (BSA) prior to incubation in primary antibody overnight in 0.5% BSA. They were then incubated in secondary antibody diluted in phosphate buffered saline prior to mounting with Vectashield 4’,6-diamidino-2-phenylindole DAPI (Vector Laboratories, Burlingame, CA). Images were then taken on the Nikon Eclipse Ti confocal microscope (Nikon USA, Melville, NY). Primary antibody used was CD68 (Invitrogen, MA5-13324) at 1:100 concentration. Secondary antibody was used at 1:250 concentration (Alexa Fluor 488 or 594 conjugated secondary 1:500 goat anti-rabbit IgG from Thermo Fisher Scientific). For Western blots, tissue was homogenized in RIPA buffer containing protease and phosphatase inhibitors. Protein was quantified by BCA assay, heated at 99 °C for 7 min to denature protein, and then run on a gel and transferred on to PVDF by wet transfer. Thereafter, samples were blocked with 5% bovine serum albumin in TBS-T, incubated in primary antibody at 1:1000 concentration overnight in blocking buffer, and then incubated with 1:5000 c concentration of HRP-conjugated secondary antibody in 5% milk in TBS-T. After washing and incubation in chemiluminescent substrate, blots were imaged on an iBright 1500 imager (Thermo Scientific). Blots were quantified using ImageJ. Primary antibodies used included IL1β (Cell Signal, D6D6T, 31202) and CD68 (Invitrogen, MA5-13324).

